# Stable Biomarker Identification For Predicting Schizophrenia in the Human Connectome

**DOI:** 10.1101/711135

**Authors:** Leonardo Gutiérrez-Gómez, Jakub Vohryzek, Benjamin Chiêm, Philipp S. Baumann, Philippe Conus, Kim Do Cuenod, Patric Hagmann, Jean-Charles Delvenne

## Abstract

Schizophrenia, as a psychiatric disorder, has recognized brain alterations both at the structural and at the functional magnetic resonance imaging level. The developing field of connectomics has attracted much attention as it allows researchers to take advantage of powerful tools of network analysis in order to study structural and functional connectivity abnormalities in schizophrenia. Many methods have been proposed to identify biomarkers in schizophrenia, focusing mainly on improving the classification performance or performing statistical comparisons between groups. However, the stability of biomarkers selection has been for long overlooked in the connectomics field. In this study, we follow a machine learning approach where the identification of biomarkers is addressed as a feature selection problem for a classification task. We perform a recursive feature elimination and support vector machines (RFE-SVM) approach to identify the most meaningful biomarkers from the structural, functional, and multi-modal connectomes of healthy controls and patients. Furthermore, the stability of the retrieved biomarkers is assessed across different subsamplings of the dataset, allowing us to identify the affected core of the pathology. Considering our technique altogether, it demonstrates a principled way to achieve both accurate and stable biomarkers while highlighting the importance of multi-modal approaches to brain pathology as they tend to reveal complementary information.

## Introduction

Schizophrenia (SZ) is a severe psychiatric disorder characterized by hallucinations and delusions, as well as impairments in memory, attention, executive and other high-order cognitive dysfunctions (1). The development of magnetic resonance imaging (MRI) has offered an effective way to examine white and grey matter changes of the brain and has motivated numerous scientists to explore the underlying neuropathology of SZ(2, 3). Over the past few years, advances in high-field structural and functional neuroimaging have made it possible to map the macroscopic neural wiring system of the human brain (4, 5) with many studies showing reproducible alterations of both structural and functional connectivity in SZ(6–9). These studies with focus on connectivity abnormalities have marked a shift in support of schizophrenia conceptualisation as a dysconnectivity syndrome (10) and thus demonstrated the importance of structural and functional connectivity in characterisation of schizophrenia(11). As such, a thorough characterisation of the structural and functional connectome is of importance to the development of novel biomarkers both for prediction and treatment (12, 13). From the connectivity standpoint multi-modal analysis has demonstrated aberrant behaviours in fronto-striatal, fronto-thalamic and fronto-temporal coupling in SZ (14) supported by studies on structural connectivity showing decreased white-matter integrity in frontal, temporal and parietal regions with studies of functional connectivity showing changes in activation in frontal and parietal areas(15) Studies from topological perspective have shown alterations of both structural and functional brain topology in SZ characterized by a less efficient global brain network organization and a limited capacity of functional integration (7). Further research using diffusion spectrum imaging has reported brain areas mainly responsible for the loss of global integration and segregation properties with prefrontal, pericentral, superior, left temporal-occipital and thalamic areas as well as striatum (7, 16, 17).

However, such findings were identified using conventional univariate strategies performing a separate statistical test at each edge of the connectome under scrutiny, thereby requiring excessively stringent corrections for multiple comparisons. On the other hand, multivariate methods are promising, although they require specialized approaches when the number of parameters dominates the observations (18). In this study, we adopt a machine learning approach that aims at discovering the most relevant set of biomarkers for discriminating subjects groups and thus quantitatively describing the group differences, both in terms of classification accuracy and stability of selected features.

### Machine learning and automatic biomarker selection

The identification of regions or connections of interest associated with a neural disorder is referred to as biomarker discovery. The identification of such biomarkers in schizophrenia could lead to clinically useful tools for establishing both diagnosis and prognosis. From a machine learning perspective, the choice of biomarkers can be addressed as a feature selection problem, aiming to find a subset of relevant features allowing us to differentiate patients from control subjects accurately.

In this work, we perform an automatic feature selection procedure in order to identify biomarkers that are relevant for the diagnosis of schizophrenia from brain connectivity data. In this context, biomarkers, therefore, correspond to structural or functional links between neural Regions of Interest (ROIs).

A key challenge in feature selection lies in the fact that diverse feature selection methods might result in different sets of retrieved features. Even when using the same technique, it may produce different results when applied to different splittings of the data. When the dimensionality of the input data is large and exceeds the number of training examples, the complexity grows by several orders of magnitude (19, 20). Therefore, the problem of deciding between two subsets of features has significant uncertainty that needs to be addressed.(21).

These issues underline the need to integrate the stability in the feature selection process so that the method can retrieve consistent features across random subsamplings of the dataset. This is especially true in a biomedical context (**?**) where many authors have focused on improving the classification performance in several mental disorders such as schizophrenia (23), Alzheimer (24), depression (25). Even though the stability of biomarker selection has been studied mainly in genomics and proteomics (21, 26), the stability of feature selection has been overlooked in the connectomics community. Therefore, we propose a general framework for stability analysis of selected features, thereby enabling the robust identification of impaired connections in the connectome of schizophrenic patients. The proposed approach is extendable to other brain disorders as well.

In the present work, we use Support Vector Machines (SVM) as a classifier (19). This is a supervised machine learning method that aims to classify data points by maximizing the margin between classes in a high-dimensional space. This classifier offers state of the art classification performances on a wide range of applications and is particularly appropriate for high-dimensional problems with few examples. The SVM classifier has been integrated into an embedded feature selection approach (27). The so-called Recursive Feature Elimination with Support Vector Machine (RFE-SVM) technique was first introduced to perform gene selection for cancer diagnosis on microarray data (28). It has been used for mapping and classification of fMRI spatial patterns on voxels (29), and functional connectivity (30). More recently, it has also been used on human brain networks to identify differences in structural connectivity related to gender (31). This method trains an SVM classifier removing the less important features and iteratively re-estimating the classifier with the remaining features until reaching the desired number of them. Accordingly, we adopt the RFE-SVM approach to automatically select brain connections that lead to the best discrimination between patients and controls, and consequently to highlight brain regions that are responsible for the disease.

The aim of the present work is threefold: First, we investigate the effect of structural, functional, and multi-modal (structural+functional) connectome with different resolutions in the classification performance of schizophrenia. Second, we perform a careful feature selection procedure across modalities in order to assess the robustness of the selected features providing the best trade-off between high accuracy and stability. Finally, the analysis of retrieved biomarkers allows us to identify a distributed set of brain regions engaged in the discrimination of patients and control subjects.

This paper is organized as follows: Section 2 introduces the properties of the dataset, the procedure for connectomes estimation as well as the general protocol we used in biomarkers identification. In section 3, we present the results on stability, classification performances, and identification of brain areas indicative of the pathology. Finally, in section 4, we lead a discussion on our findings and conclusions.

## Materials and methods

### Subjects

For this study, two age-balanced groups were considered. The cohort consisted of a schizophrenic group of 27 subjects with a mean age of 41.9 ± 9.6 years and a control group of 27 healthy subjects with a mean age of 35 ± 6.8 years. The patients in the schizophrenic group were recruited from the Service of General Psychiatry at the Lausanne University Hospital. They met DSM-IV criteria for schizophrenic and schizoaffective disorders (American psychiatry association, 2000^1^). Healthy controls were recruited throught advertisement and assessed with the Diagnostic Interview for Genetic Studies (32). Subjects with major mood, psychotic, or substance-use disorders and having first-degree relative with a psychotic disorder were excluded. Moreover, a history of neurological disease was an exclusion criterion for all subjects. This study was carried out with 24 out of the 27 schizophrenic patients being under medication with chlorpromazine equivalent dose (CPZ) (average medication 431 ± 288 mg)(33). We obtained written consent from all the subjects following the institutional guidelines approved by the Ethics Committee of Clinical Research of the Faculty of Biology and Medicine, University of Lausanne, Switzerland. The dataset is public and available in Zenodo platform^2^ (34).

### Brain network estimation Magnetic resonance imaging

All subjects were scanned on the 3 Tesla Siemens Trio scanner with a 32-channel head coil. Three acquisition protocols were part of the MRI session, namely structural, functional and diffusion MRI scans. Structural MRI: magnetization-prepared rapid acquisition gradient echo (MPRAGE) sequence with in-plane resolution of 1 mm, slice thickness of 1.2 mm of total voxel number of 240×257×160 and TR, TE and TI were 2300, 2.98 and 900 ms respectively. Diffusion MRI: diffusion spectrum imaging (DSI) sequence with 128 diffusion-weighted images of b0 as a reference image and a maximum b-value of 8000 s/mm2. The time of acquisition was 13 min and 27s. The number of voxels was 96×96×34 with a resolution of 2.2×2.2×3.0 mm, and TR and TE were 6100 and 144 ms respectively. The issue of motion-artifacts linked to signal drop-outs was dealt with by visually inspecting the signal, and no subject had to be excluded as a result of this (35). Functional MRI: a resting-state functional MRI (fMRI) acquired for 8 minutes (3.3×3.3×3.3 mm voxel size, TR = 1920ms, TE = 30ms, 32 slices, flip angle 85°). During the fMRI acquisition, subjects were asked not to fall asleep and let their mind wander while fixating their vision to the cross on the screen.

### Structural networks

Structural and diffusion MRI data were used to estimate the weighted and undirected structural connectivity matrices in the Connectome Mapping Toolkit (16, 36, 37). Firstly, white matter, grey matter, and cerebrospinal fluid segmentation was performed on the structural data and further linearly registered to the b0 volumes of the DSI dataset. Secondly, the first three scales of the Lausanne multi-scale atlas were used to parcellate the grey matter. In detail, the first scale consisted of 68 cortical brain regions and 14 subcortical regions with scale two and three subdividing the first scale into 114 and 219 cortical regions (37). Further, deterministic streamline tractography, estimating 32 diffusion directions per voxels, was used to reconstruct the structural connectivities from the DSI data(38). The normalized connection density quantified the structural connectivity between brain regions and is defined as follows,

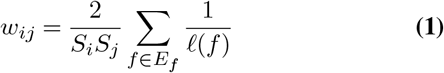

where *w* represents an edge between brain regions *i* and *j*, *S_i_* and *S_j_* are the surface areas of regions *i* and *j*, *f* ∈ *E_f_* represents a streamline *f* in the set of streamlines *E* and *ℓ*(*f*) is a length of a given streamline *f* (4, 16). The normalisation by brain region surfaces accounts for their slightly varying size and the streamline length normalisation accounts for a bias towards longer connections imposed by the tractography algorithm.

### Functional Networks

Functional connectivity matrices were computed from fMRI BOLD time-series. Firstly, the first four-time points were excluded, yielding the number of time points to be *T* = 276 (39). Rigid-body registration was applied to individual timeslices for motion-correction. The signal was then linearly detrended and corrected for physiological confounds and further motion artifacts by regressing white-matter, cerebrospinal fluid, and six motion (translations and rotations) signals. Lastly, the signal was spatially smoothed and bandpass-filtered between 0.01 — 0.1 Hz with Hamming windowed sinc FIR filter. Linear registration was performed between the average fMRI and MPRAGE images to obtain the ROIs timeseries(40). An average timecourse for each brain region was computed for the three atlas scales. In order to obtain the functional matrices, the absolute value of the Pearson correlation was computed between individual brain regions’ timecourses. All of the above was performed in subject native space with Connectome Mapper Toolkit and personalized Python and Matlab scripts(36)(41).

### Biomarker evaluation protocol

Our evaluation methodology is based on Abeel et al 2010 (26) used for biomarker identification in cancer diagnosis on microarray data. In order to assess the robustness of the biomarker selection process, we generate slight variations of the dataset and compare the outcome of selected features across these variations. Therefore, for a stable marker selection algorithm, small variations in the training set should not produce important changes in the retrieved set of features. We perform a nested 5-fold cross-validation (CV) approach. Here, the external CV is used to provide an unbiased estimate of the performance of the method, whereas the inner CV loop is used to fitting, tunning and selecting the optimal parameters of the model. Concretely, we generate 100 subsamplings of the original dataset, shuffling the outer 5-fold CV scheme 20 times. The 80% of the data, i.e., four folds of the outer CV (pink color in Figure 1), is used as training set within the inner CV, where the best model and features are selected. That is, four folds are used as training set and the held-out fold as validation set to tune the parameters of the model. The model achieving the best performance on the validation set is selected together with the features selected by the RFE-SVM method. The remaining 20% of the outer CV, i.e., the hold-out fold, is used as testing set to provide an unbiased evaluation of the final model and assess the performance of the classifier. Therefore, the overall accuracy is given by the average testing accuracy across subsamplings. See Figure 1 for a schematic view of the methodology.

**Fig. 1.**
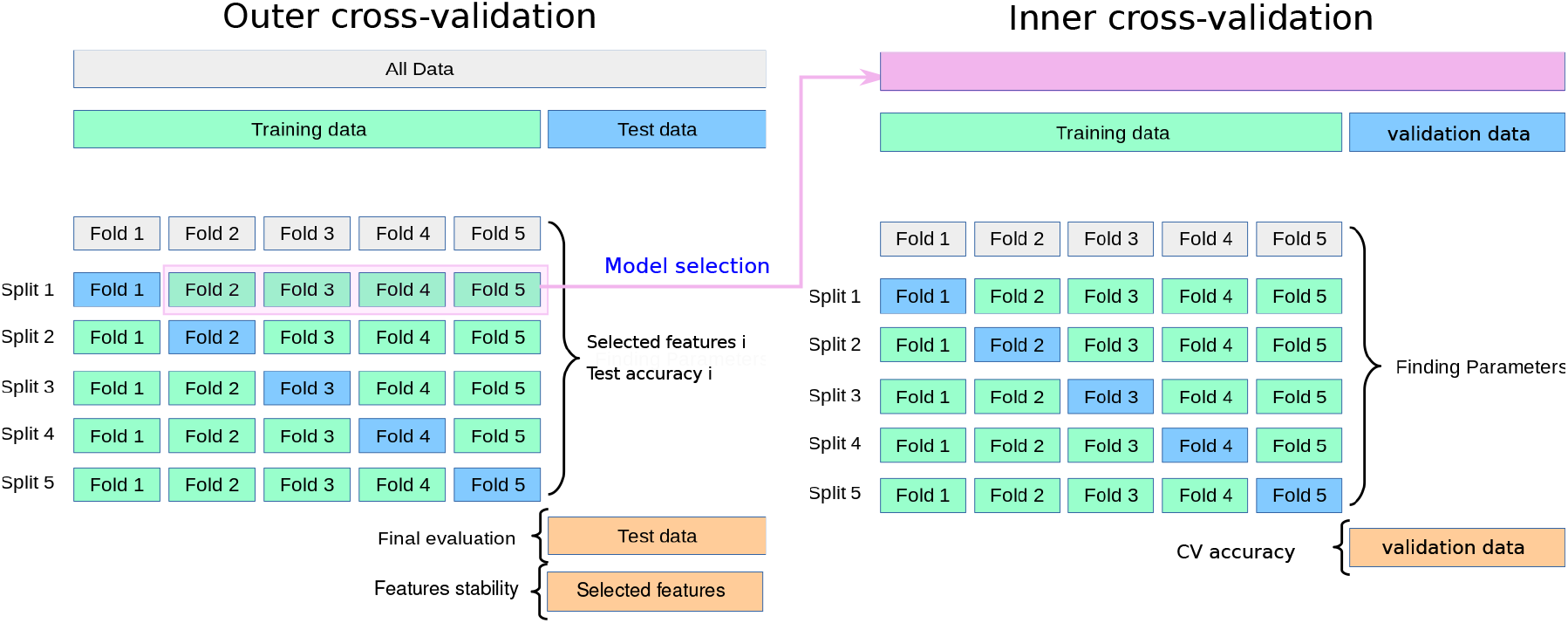
Overview of the proposed method. The figure represents the nested 5-fold CV subsampling of the entire dataset, i.e., top-left gray bar. (Left) The outer CV is used to evaluate the performance of the model. The 80% of the data, i.e., four folds (pink box), is used as training set, where the best model and features are selected. The remaining 20% is used as testing set, to evaluate the performance of the model. (Right) Within the inner CV, four folds are used for training and the hold-out fold as validation set. The best model, features and parameters are selected according with the best CV accuracy. The outer CV is shuffled 20 times, generating 100 subsamplings of the dataset and therefore the same number of selected features ‘fingerprints’. The stability of selected biomarkers and the final accuracy is assessed over all subsampling estimations.

### Embedded feature selection (RFE-SVM)

In this work we use a linear SVM classifier (19). SVM has proven state of the art performance in computational biology (42) in particular with problems of very high dimension, scaling very well as a function of the number of examples. Given a set of data examples ***x**_i_* ∈ ℝ^*p*^, *i* = 1,…,*n*, and a vector ***y*** ∈ {1, −1}^*n*^” representing the group membership of data points, SVM aims to find the hyperplane that has the largest distance to the nearest training data points of any class. The mathematical formulation can be written as an optimization problem in its primal form (19):

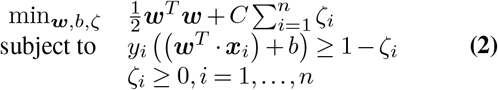

where *ζ_i_* are slack variables controlling the overlapping between classes cased by noisy examples, and *w* ∈ ℝ^*p*^ is a normal vector of the hyperplane. A classifier which generalizes well is then found by controlling both the classifier capacity ||*w*||, and the sum of the slacks 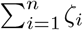. (19). The solution of the optimization problem in its dual form provides the coefficients of such a hyperplane as the sum of the support vectors *α_i_*, i.e., points lying on the max-margin hyperplane of separation between classes, and the training examples ***x**_i_* as:

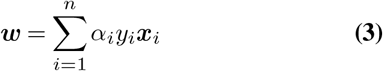

These coefficients can be interpreted as a strength or contribution of each feature to the decision of the hyperplane. As a consequence, the square value of each coefficient (or weight) can be used as a score to rank features from the most to the least important for the selection process.

Recursive feature elimination SVM (RFE-SVM) (28) is an iterative algorithm integrating a ranking criterion for eliminating features in a backward fashion. Starting with the whole set of features, a linear SVM is estimated using the training set, and their features are ranked according to the weights assigned by the algorithm. Consequently, the least important features according with the mentioned criterion are removed and the remaining ones are used to train a new model, repeating the process until reaching a desired minimal subset of features.

The RFE-SVM algorithm has a set of internal parameters influencing the computational complexity and the accuracy of the method. The fraction *E* of features to remove at each step of RFE (also called step size) is critical for the running time. Dropping one feature at a time allows a finer selection but with a prohibitive computational cost. Setting the step size to 100%, RFE reduces to a single SVM estimation which ranks all the features in one step. Following the work of Abeel et al. (26), we drop 20% of the least relevant features at each iteration by default. Additional sensitivity analysis is reported in our experiments to check the influence of this parameter, when varies between 20,50 and 100 percent. Yet, a stopping criterion is needed to finish the iterative process. Thus, in our experiments, we dropped features until reaching a minimum of *s* ∈ {0.5,1,2,5,10,25,50} percentage of selected features (stopping criterion).

Another critical parameter is the regularization constant *C* of the SVM (Eq. 2). The *C* parameter controls the misclassification rate of the classifier. A larger value makes the optimization choose a smaller margin hyperplane, losing generalization capabilities. The smaller the *C*, the larger the margin of separation, yielding more misclassified points. Therefore this parameter influences the classification accuracy of the model. We cross-validate the optimal *C* by performing a grid-search over the set of values {0.01,0.1,1,10,100} using only the training set, which corresponds to the 80% of folds from the outer CV scheme. Finally, we introduce a random feature selector for each subsampling of the dataset where one subset of features is selected uniformly at random before training the classifier. We used Python 2.7 for implementing our approach, with the RFE-SVM implementation of scikit-learn 0.22.2.^3^

### Measuring the classification performance

Because our dataset is class balanced and the task is a binary classification problem, we adopt the accuracy as the metric to quantify the performance of classification. This metric is defined as the ratio of correct classifications to the number of classifications done as follows:

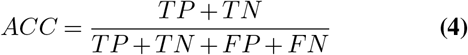

where TP is true positive (number of control subjects classified correctly), TN is the true negative (patients classified correctly), FP is false positives (number of patients classified as control subjects) and FN is false negatives (number of control subjects misclassified).

### Assessing the stability of feature selection

We consider the vectorized connectivity matrices of the connectomes as input features for the biomarker selection process. Therefore, for a given connectome, one structural feature refers to the normalized connection density between two linked brain regions, whereas a functional feature refers to the Pearson correlation between two individual brain time courses. The number of input features used for each modality and resolution is shown in Table 1.

**Table 1.** Number of input features for each modality and resolution

| Resolution | 83 | 129 | 234 |
| --- | --- | --- | --- |
| Num of features (SC/FC) | 3403 | 8256 | 27261 |
| Num of features (MM) | 6806 | 16512 | 54522 |

We consider a dataset with *M* = 54 subjects and *N* features without considering self-loops, i.e., discarding the diagonal non-zero values of the connectivity matrices. If we denote the considered connectome resolution as *d* ∈ {83,129,234}, the number of input features is 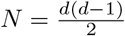 because of the symmetry of the connectivity matrices. Drawing *k* =100 subsamplings and after applying a feature selection procedure (RFE-SVM) in the 80% of each subsampling, we obtain a respective feature signature, i.e. sequence of indices of selected features. Considering two signatures *f_i_* and *f_j_* obtained from different subsamplings *i* and *j*, the stability index (43) between *f_i_* and *f_j_* is defined as:

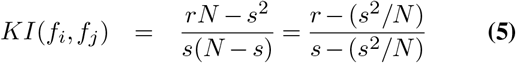

where *r* = |*f_i_* ⋂ *f_j_* | and *s* = |*f_i_* | = |*f_j_*|, the size of the signature, is the number of selected features. This quantity measures the consistency between pairs of features. For *s* and *N* fixed, *KI* increases until reaching maximum at 1 when the two subsets are identical. The minimum value is bounded from below by —1 when the subsets are perfectly disjoint and the signature size of *N*/2. The overall stability index for a sequence of signatures is defined as the average of all pairwise stability indices on *k* subsamplings:

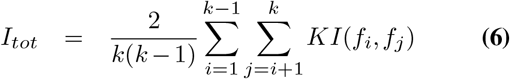

Given that *I_tot_* is bounded between —1 and 1, the greater this value the better the agreement between the selected subsets of features. In particular, a negative value for *I_tot_* indicates that the potential agreement between the selected biomarkers is mostly due to chance. In the sequel we will refer to the overall stability *I_tot_* as the Kuncheva index (43).

## RESULTS

### Connectomes classification and features stability

First, we investigated the effect of different brain connectivity modalities and different scales in the discrimination of patients and normal controls. For each case, we control the step size and the percentage of selected features of the RFE-SVM algorithm, assessing their impact on the classification accuracy and the stability of the selected features.

Figures 2, 4 and 5 show the average classification accuracy after performing RFE-SVM as well as the stability of selected biomarkers across modalities and scales. It can be seen that across scales, the functional connectivity matrices (Fig. 4) achieve better accuracies than the structural features (Fig. 2), but conversely, structural matrices are more stable than functional. However, when combining the two modalities, i.e., by concatenating features of both modalities and letting the algorithm choose a blend of structural and functional features we achieve the best performances (Fig. 5) in terms of both accuracy and stability. Note that in all figures, the curves are highly overlapping each other showing that the number of dropped features in the RFE-SVM algorithm, i.e., the step size, is not a critical parameter in this dataset. It is to be noted that in the multi-modal case, the percentage of finally selected features is divided by two since we combine twice as many features as in the case of structural or functional connectivity alone. Table 2 shows the actual number of selected biomarkers across modalities and resolutions.

**Fig. 2.**
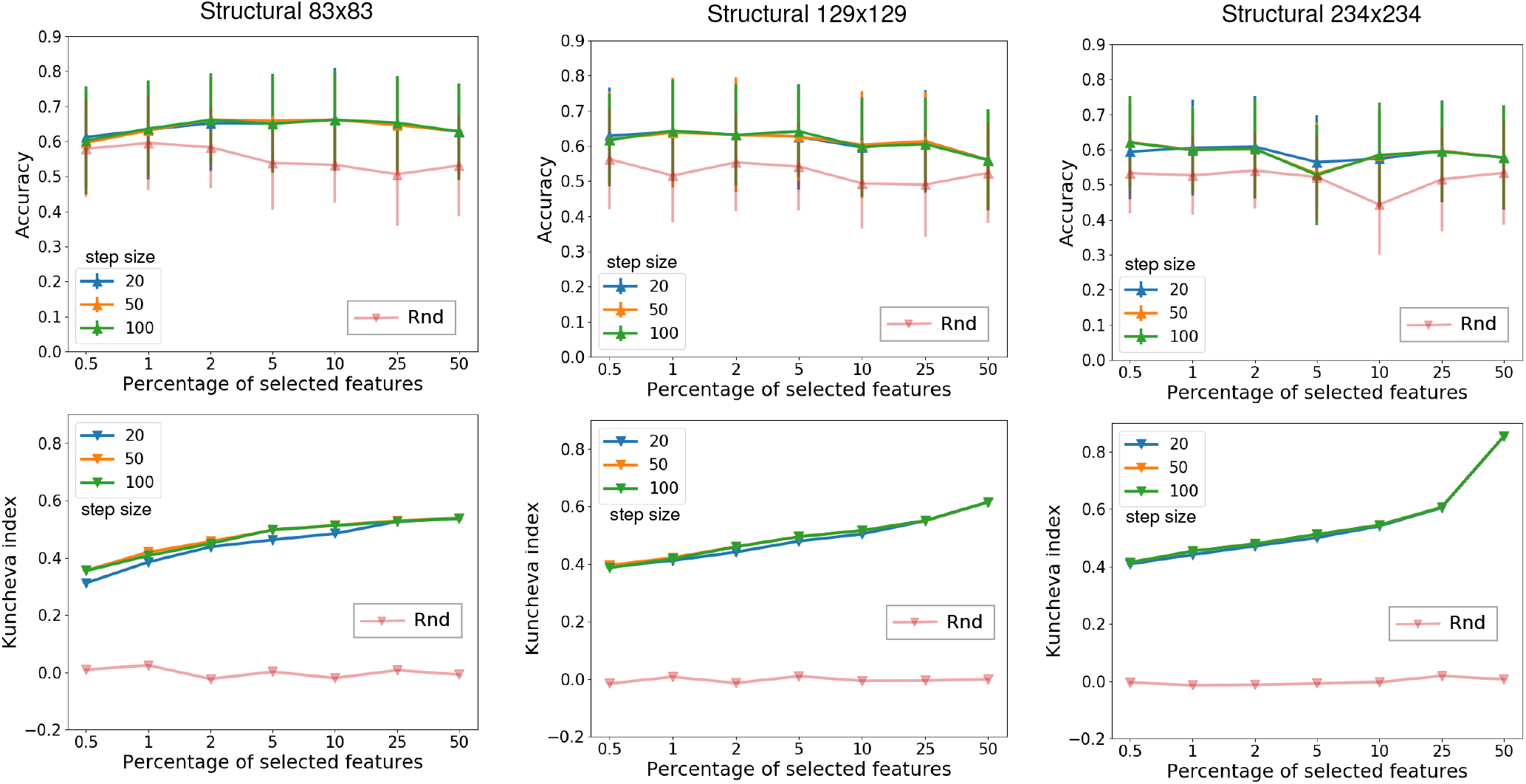
Each column represents the mean accuracy of the outer CV folds +/− standard deviation (top) and stability (bottom) for a given scale of the **structural** connectome. The step size corresponds to the percentage of features dropped at each step of the RFE-SVM algorithm. Red curve represents a random selection of features.

**Table 2.**
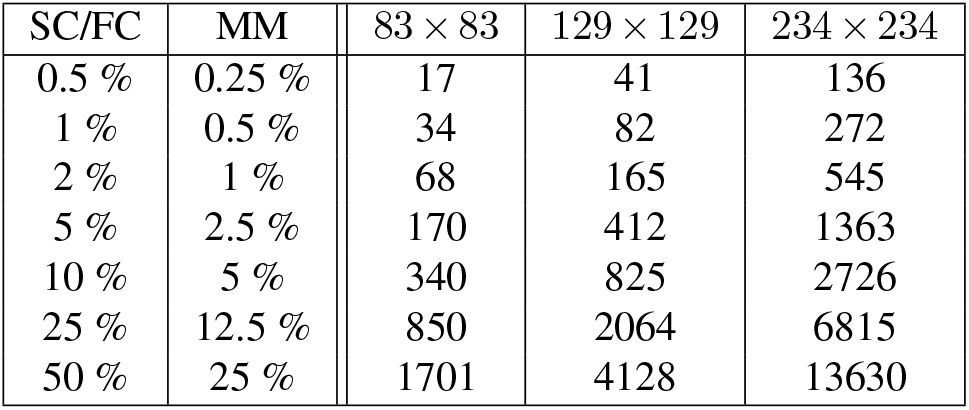
Number of selected features for SC, FC and MM connectomes

| SC/FC | MM | 83 × 83 | 129 × 129 | 234 × 234 |
| --- | --- | --- | --- | --- |
| 0.5 % | 0.25 % | 17 | 41 | 136 |
| 1 % | 0.5 % | 34 | 82 | 272 |
| 2 % | 1 % | 68 | 165 | 545 |
| 5 % | 2.5 % | 170 | 412 | 1363 |
| 10 % | 5 % | 340 | 825 | 2726 |
| 25 % | 12.5 % | 850 | 2064 | 6815 |
| 50 % | 25 % | 1701 | 4128 | 13630 |

We observe in all cases that the stability increases with the percentage of selected features, which is expected since the overlapping between signatures in Eq. 6 is more likely when more features are considered.

To further investigate the effect of modalities and resolutions on both classification accuracy and stability, we select the best scores, i.e., best accuracy with the associated stability, from Figs. 2, 4 and 5, and plot them in Fig. 3. As can be seen in Figure 3, the 234×234 structural resolution has the lowest accuracy and stability, which can be explained by the fact that as the ROIs are smaller, it makes them more susceptible to noise introduced from differences in quality of alignment of the parcellation to the native scan across the subjects. Also, the fact of averaging them over less voxels make the signal more variable between subjects, affecting the stability of the method.

**Fig. 3.**
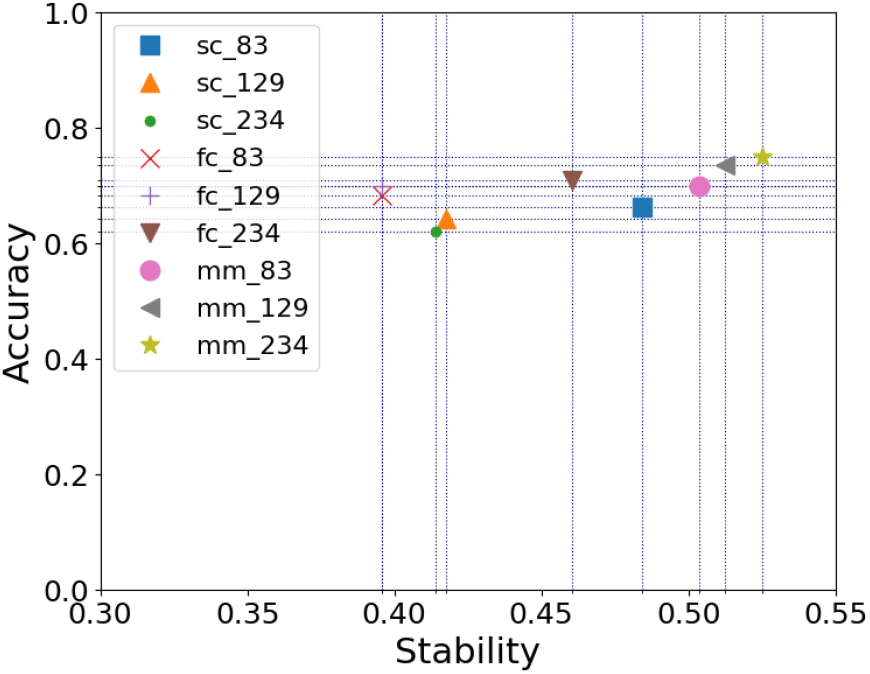
Best accuracy versus stability for all considered modalities and resolutions.

**Fig. 4.**
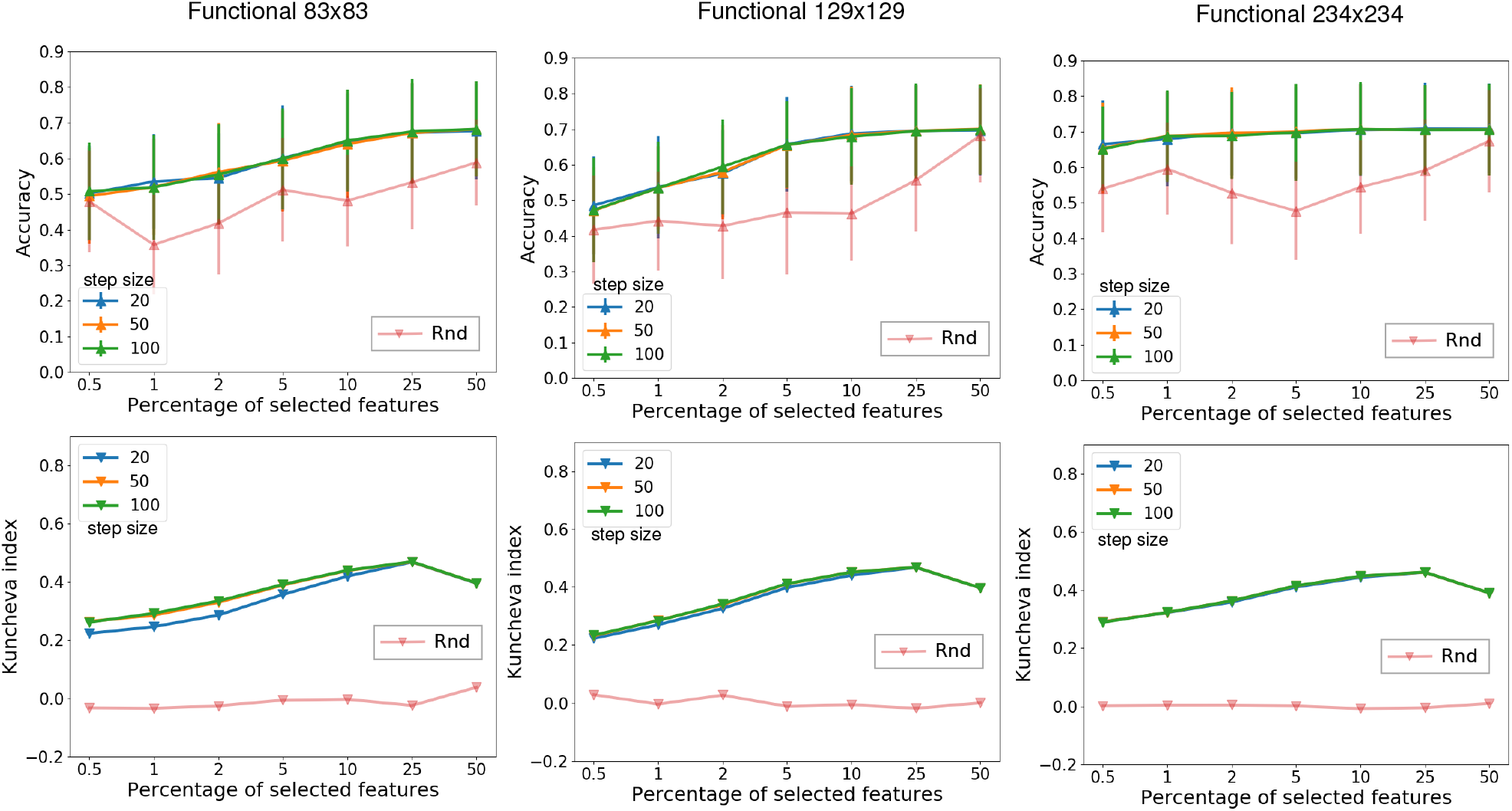
Each column represents the mean accuracy of the outer CV folds +/− standard deviation (top) and stability (bottom) for a given scale of the **functional** connectome. The step size corresponds to the percentage of features dropped at each step of the RFE-SVM algorithm. Red curve represents a random selection of features.

**Fig. 5.**
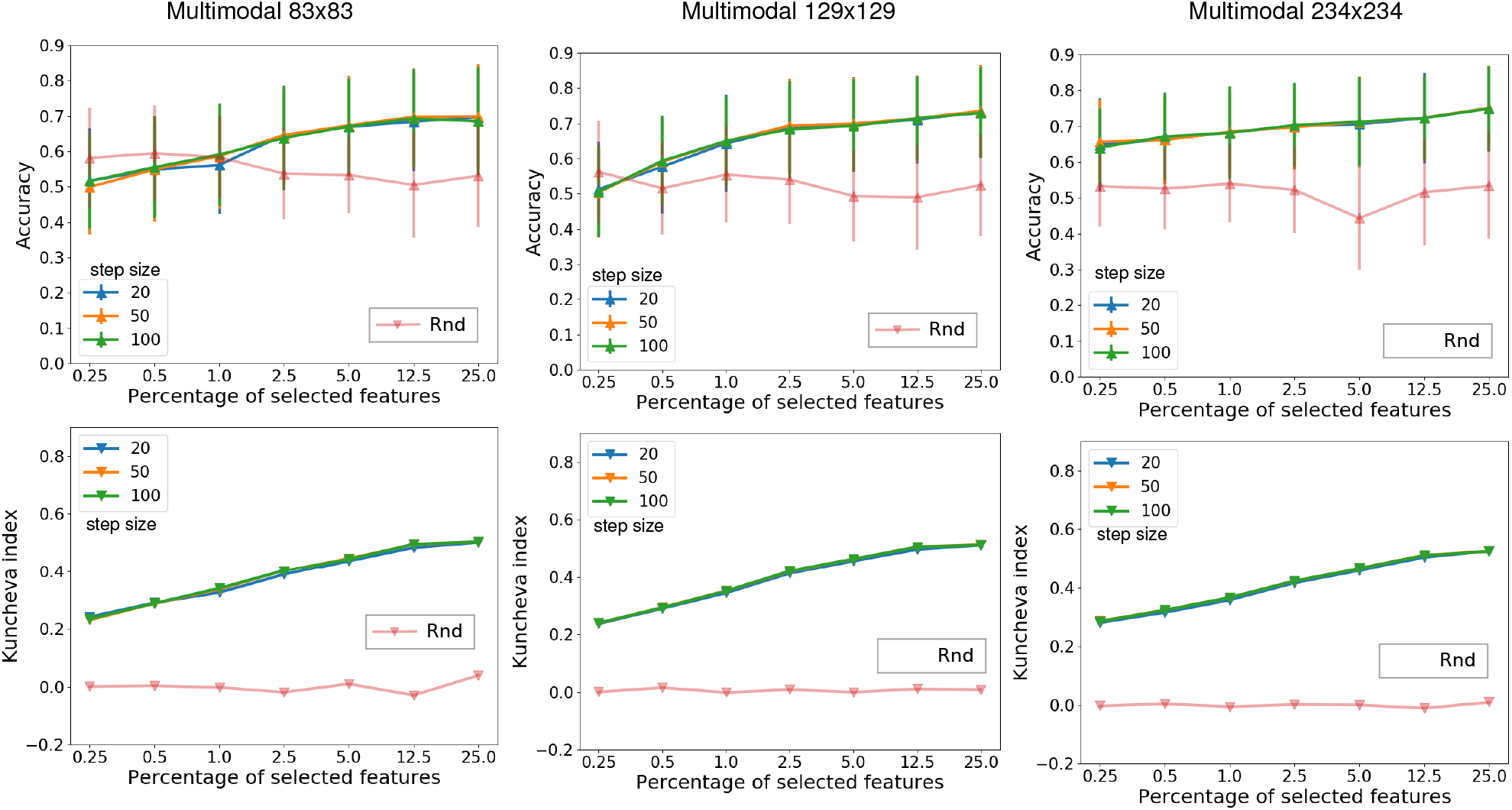
Each column represents the mean accuracy of the outer CV folds +/− standard deviation (top) and stability (bottom) for a given scale of the **multimodal** connectome. The step size corresponds to the percentage of features dropped at each step of the RFE-SVM algorithm. Red curve represents a random selection of features.

In order to quantify the respective involvement of features extracted from both modalities, we provide in Figure (6) a summarized view of the share of structural features in the total number of multimodal features. We observe that our method always selects more functional features than structural ones. This is consistent with the better accuracies obtained with functional features, and with the fact that the RFE-SVM ultimately optimizes such accuracy scores. However, the share of selected structural features is always non-zero (although it decreases with the connectome resolution), thereby indicating their respective importance in the retrieved features sets. It is interesting to observe that structural and functional features alone achieve the best stability and accuracy respectively, whereas the multimodal combination achieves the best trade-off between both metrics, for all resolutions.

**Fig. 6.**
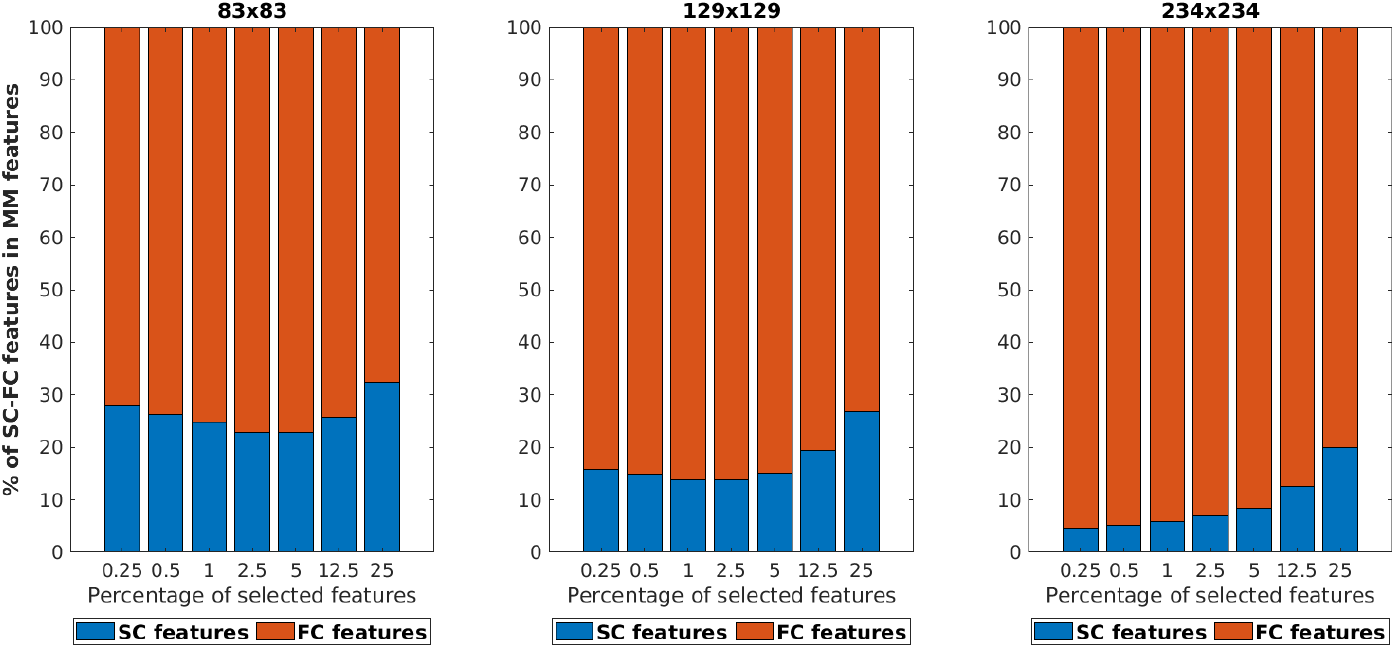
Percentage structural and functional features selected from the multi-modal connectome across resolutions.

### Identification of brain regions in schizophrenia diagnosis

We proceed with the identification of brain areas involved in the classification of patients and controls. For simplicity in the identification of brain regions and comparison with other authors, we analyze the results for the multi-modal 83 × 83 connectome, but we reported the results with other resolutions in the appendix section.

Selected features in the graph space correspond to links representing either connection densities in the structural matrices or Pearson correlations in the functional connectome, see Figure 7. Furthermore, inspecting the frequency of each selected feature across subsamplings informs us about the overall relevance of the edge in the classification. In other words, the frequency of an edge is indicative of the importance of the associated ROIs in the classification task.

**Fig. 7.**
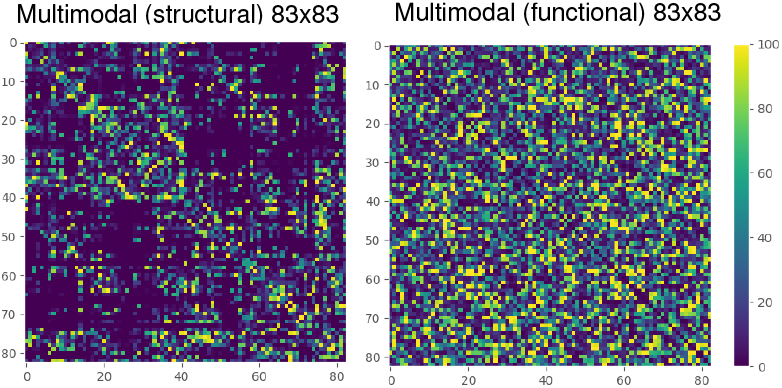
Selected markers in the multimodal 83 × 83 connectome. Colormap shows the number of times each feature is selected across subsamplings

Given a brain connectivity matrix at a resolution *r* we define *W* as the *r* × *r* matrix where the element *w_i,j_* encodes the frequency at which the edge (*i,j*) is selected as relevant across subsamplings (see Fig. 7). Thus, the *degree of relevance* of an ROI *i* reads:

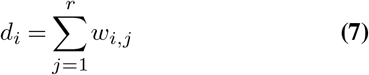

Figures 8 and 9 represent the degree of relevance of brain regions for the multi-modal 83 × 83 connectome (structural and functional respectively), sorted in decreasing order. We defined the affected core (a-core) to be composed of brain areas with a degree of relevance higher than the overall average. As can be seen in Fig. 8, our affected core overlaps the a-core definition of Griffa et al. 2015 (16) (shown as blue bars). It can be notice that while caudalmiddle-frontal, inferiortemporal, postcentral, precentral, parsoper-cularis and parsorbilatis regions are included in the Griffa’s a-core, our method does not consider them assigning them a lower degree of relevance. A similar behavior can be seen at the right-hemisphere where our method discards Griffa’s a-core regions such as caudalmiddlefrontal, medi-alorbitofrontal, parstriangularis, postcentral, rostralmiddle-frontal and supramarginal.

**Fig. 8.**
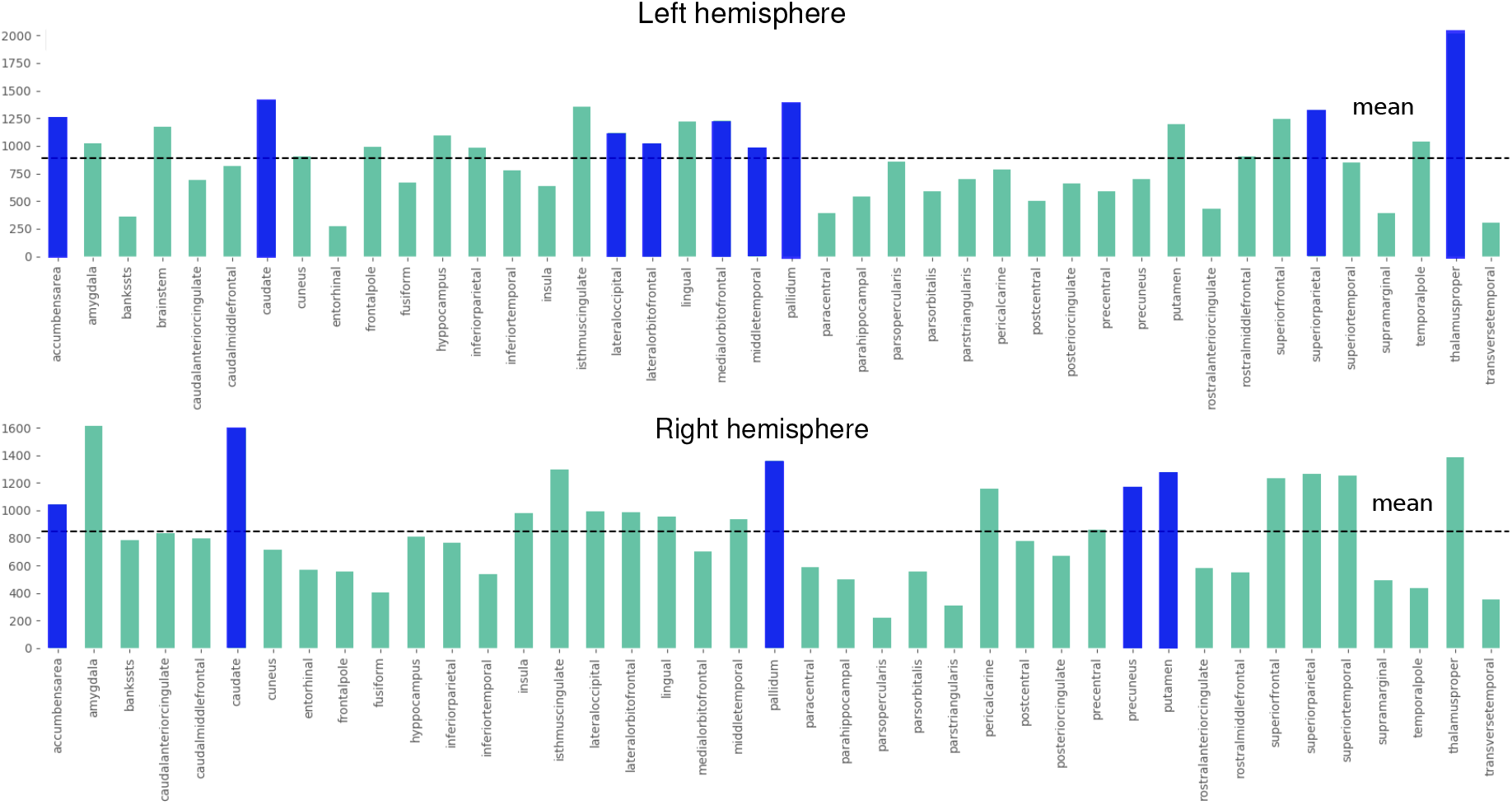
Degree of relevance for ROIs in the structural mode of the multimodal 83 × 83 connectome. Blue bars correspond to Griffa’s a-core overlapping. The horizontal line is the average degree of relevance. ROIs above the mean belong to our a-core.

**Fig. 9.**
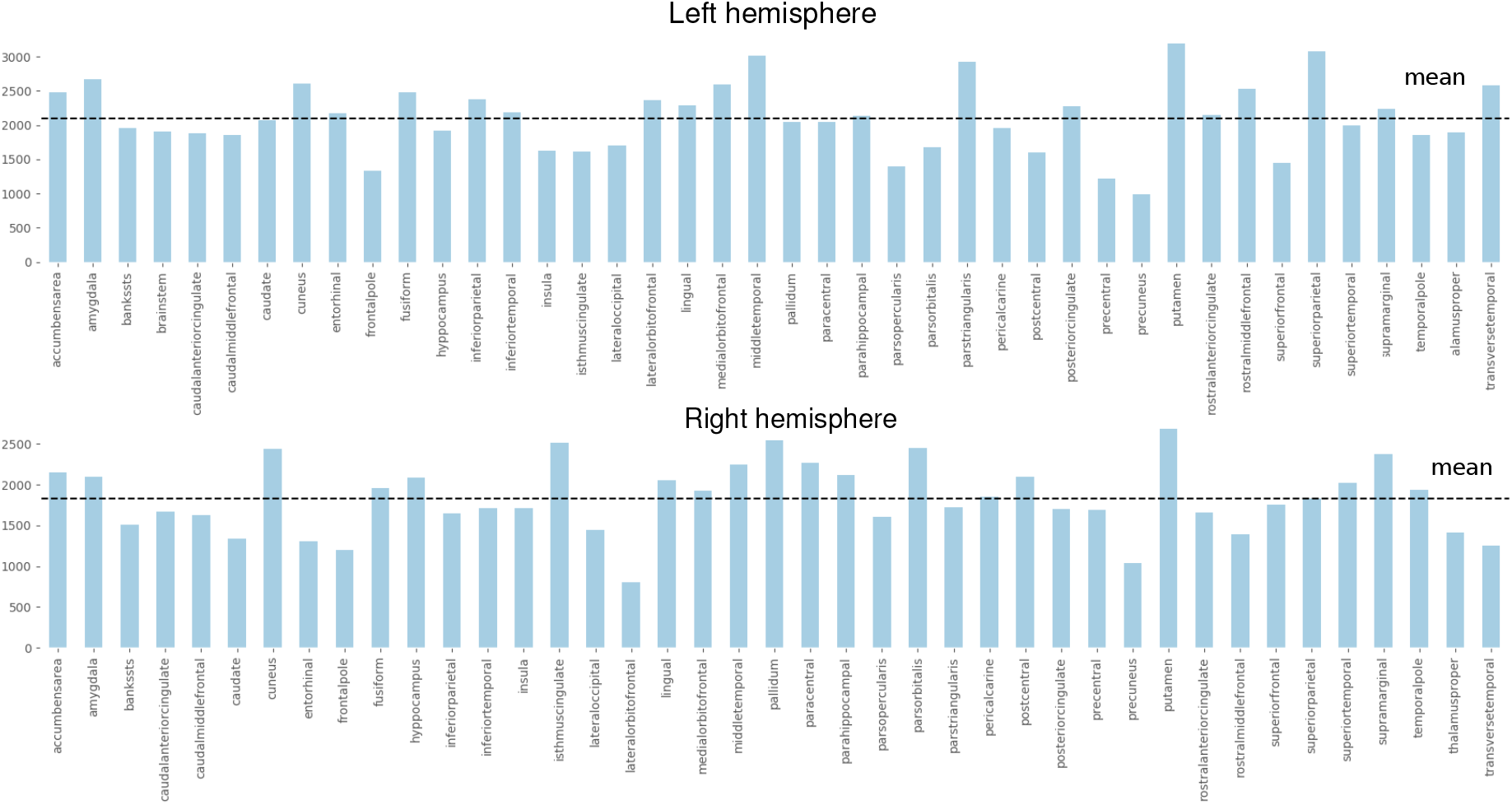
Degree of relevance for ROIs in the functional mode of the multimodal 83 × 83 connectome. The horizontal line is the average degree of relevance. ROIs above the mean belong to our a-core.

We plot the brain surface in Figure 10, 11, normalizing both SC and FC by the sum of all their connections and plotted the regions above the mean distribution.

**Fig. 10.**
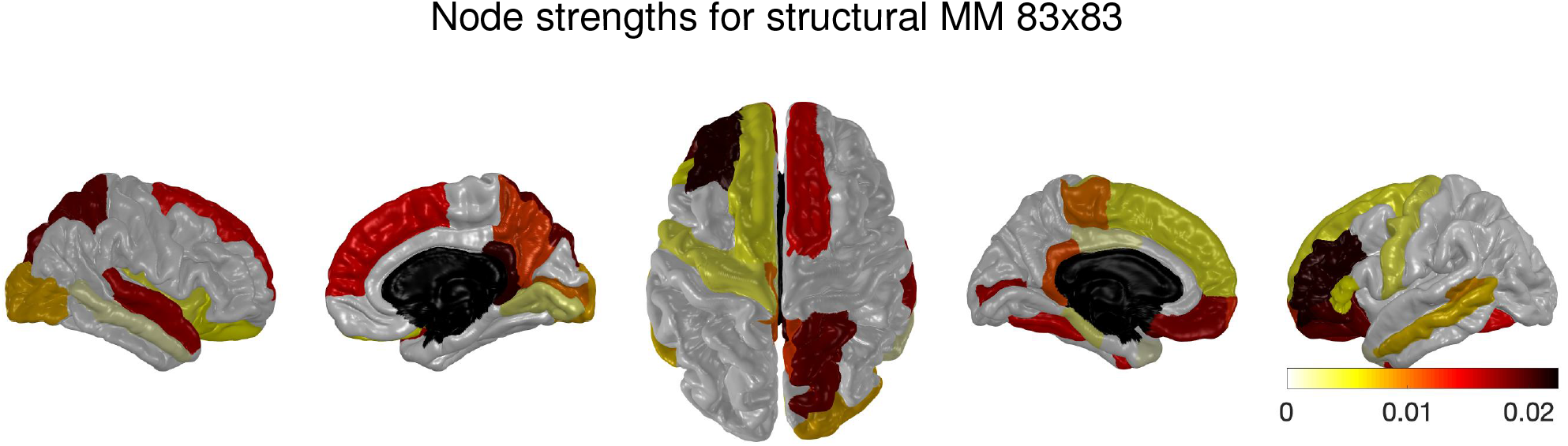
Brain surface representation of brain areas with higher relevance degree than the average for the 83 × 83 resolution of the multi-modal structural mode.

**Fig. 11.**
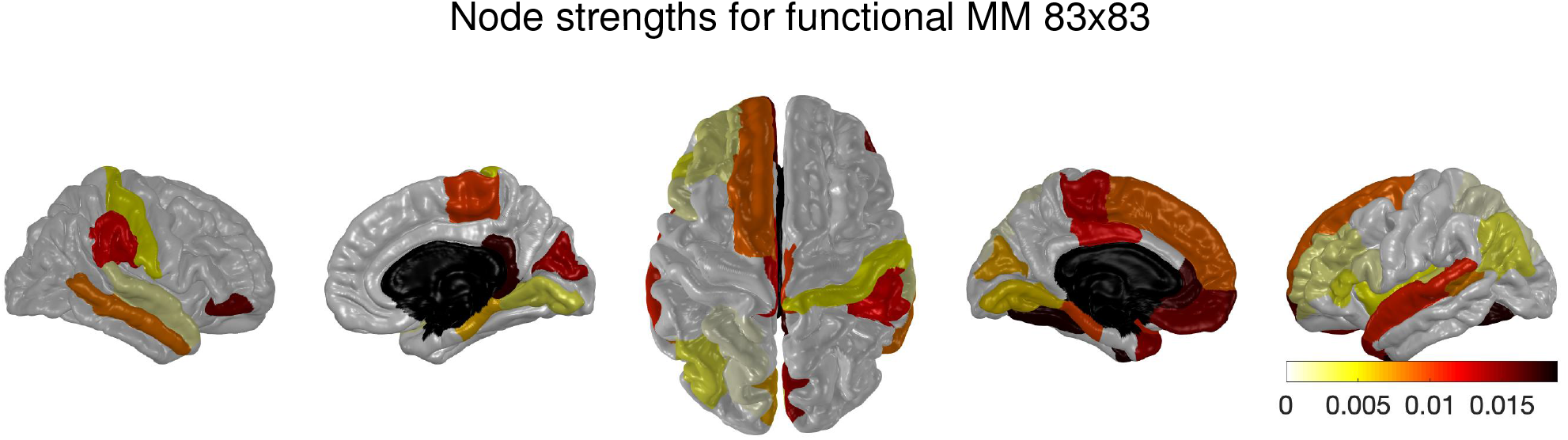
Brain surface representation of brain areas with higher relevance degree than the average for the 83 × 83 resolution of the multi-modal functional mode.

## DISCUSSION

This paper has investigated the effect of different connectivity modes of the human connectome in the selection of robust biomarkers for the identification of subjects with schizophrenia. We perform an automatic feature selection process on the edge space aiming to retrieve a compact subset of meaningful biomarkers performing accurately on the identification of schizophrenia versus healthy controls. Besides, we analyze the robustness of the retrieved features concerning the sample variation, based on the fact that stable biomarkers will not change dramatically in different subsamplings of the underlying dataset (26).

We found out that combining structural and functional connectivity matrices as a multi-modal representation of connectomes provides the best trade-off between high accurate and stable biomarkers in schizophrenia classification. As such, we identified the affected core of the pathology from the retrieved features and carried out a confirmatory analysis of the structural alterations of this core showing decrease both in gFA and iADC of edges within the a-core with no change for edges outside of a-core in patients compared to healthy subjects.

It is well known that anatomical connections constrain functional communications (44). Structural and functional connections have been shown to correlate(4) and white-matter pathways have accurately predicted functional connectivity (45). Thus, both modalities can yield a very unique information about the pathology. The fact that the best accuracy and stability was achieved by combining SC and FC features reflects that the two modalities are not well coupled (46). Furthermore, it highlights the importance of understanding both approaches in the explanation of complex neural disorders in terms of structure and function, at least insofar the prediction of them is concerned.

Based on the frequency from which the RFE-SVM algorithm selects relevant edges, we can map edges to the node space by looking at the strengths of connection densities or correlation between brain regions. The degree of relevance for nodes allows us to identify the affected core as the brain regions that are highly active during the training phase and therefore, are selected more often for relevant edges. Our findings slightly overlap results from Griffa et al. 2015 (16) and provide further evidence of brain regions involved in the pathology. Furthermore, they offer a unique perspective in defining the affected core both from the influence of SC and FC and hence defining an SC-FC a-core which might give a richer description of the regions that are affected in SCHZ (8, 16).

Finally, it is worth mentioning that this work was performed on a small sample dataset which is not suited to identify regions with great confidence. Here we present a proof of concept having in mind the importance of using it in larger dataset and especially in subjects with At Risk Mental State (ARMS) who have not yet developed a full blown illness.

## CONCLUSION

In summary, we investigated the classification performance between schizophrenic patients and healthy subjects in the structural, functional and multimodal connectomes of varying atlas resolutions. Moreover, we focused on robustness of the selected features and thus enriched the outcome of the classification to the trade-off between high accuracy and stability. We showed that a combination of structural and functional connections achieves the best performance possibly due to the complementary nature of the information content both modalities hold. Lastly, we used the biologically relevant features to define the affected core of the pathology and confirmed the structural alterations associated with the affected core edges between schizophrenic patients and healthy controls. By providing an important addition to the classification of human pathologies in form of stability analysis, we hope this study to be a step in the right direction for the diagnosis and treatment in the clinical practice.

## Acknowledgments

This work was supported by Concerted Research Action (ARC) supported by the Federation Wallonia-Brussels Contract ARC 14/19-060, Flagship European Research Area Network (FLAG-ERA) Joint Transnational Call “FuturICT 2.0”; Swiss National Science Foundation grant numbers: 156874 and 158776 to which are gratefully acknowledged. Benjamin Chiêm is a FRIA (F.R.S.-FNRS) fellow.

## A-core for other parcelations

In this appendix we show the identification of brain regions for the multi-modal 129 × 129 and 234 × 234 connectomes.

**Fig. 12.**
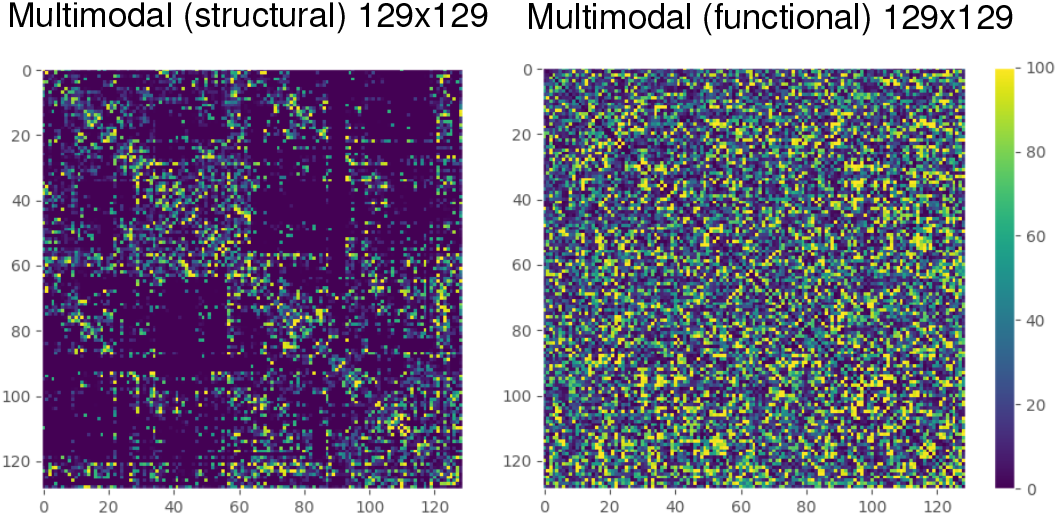
Selected markers in the multimodal 129 × 129 connectome. Colormap shows the number of times each feature is selected across subsamplings

**Fig. 13.**
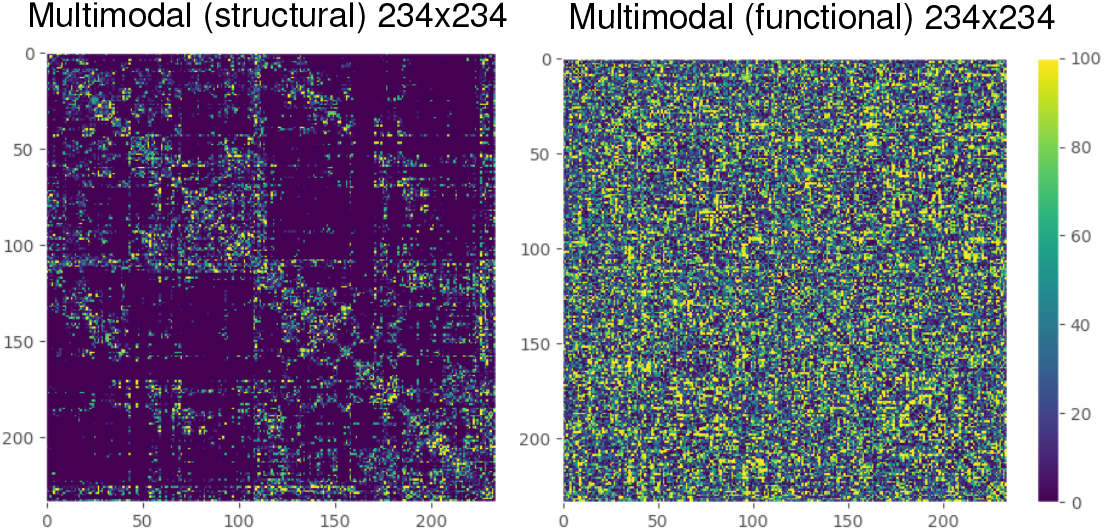
Selected markers in the multimodal 234 × 234 connectome. Colormap shows the number of times each feature is selected across subsamplings

**Fig. 14.**
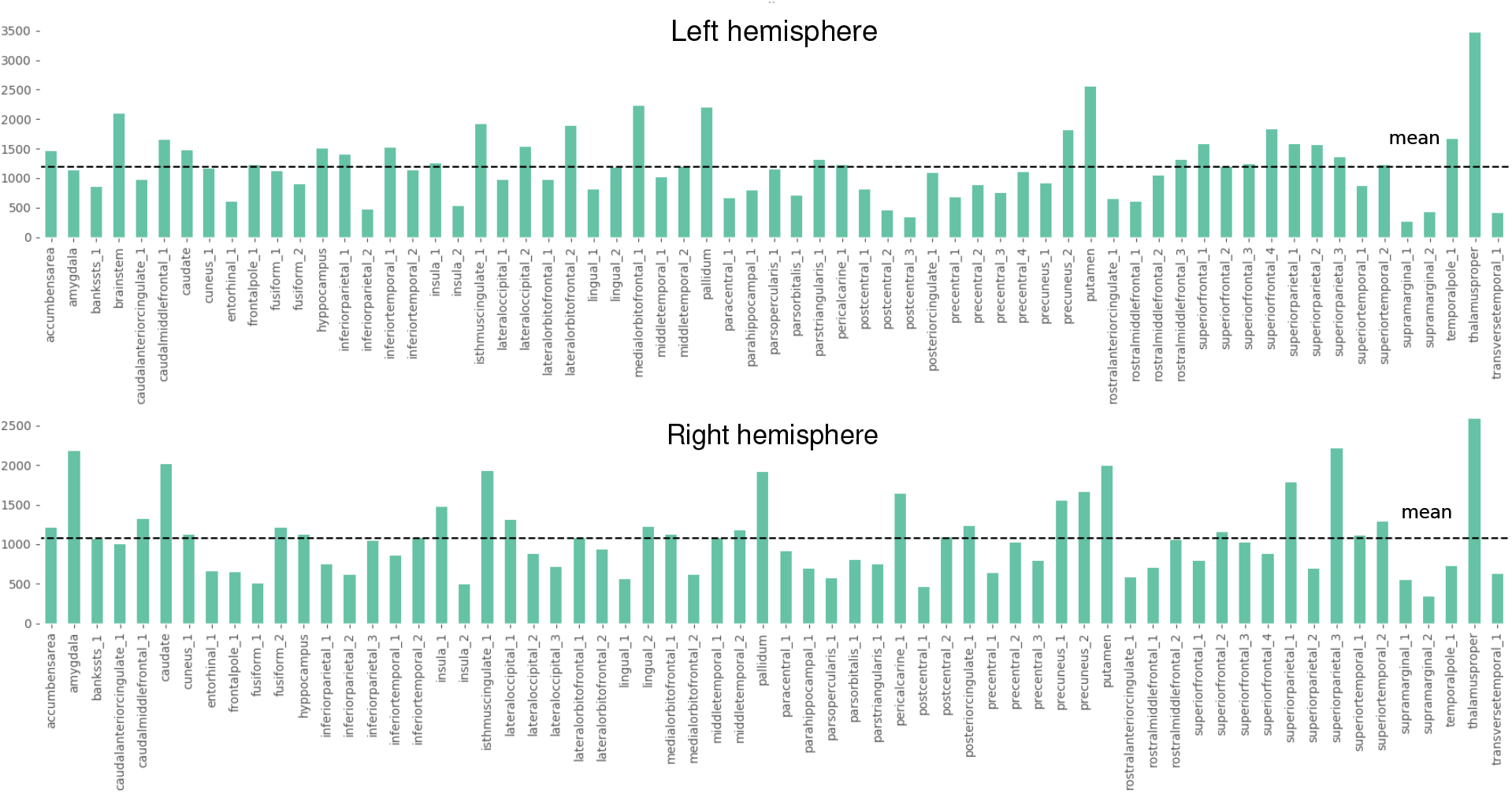
Degree of relevance for ROIs in the **structural** mode of the multimodal 129 × 129 connectome. The horizontal line is the average degree of relevance. ROIs above the mean belong to our a-core.

**Fig. 15.**
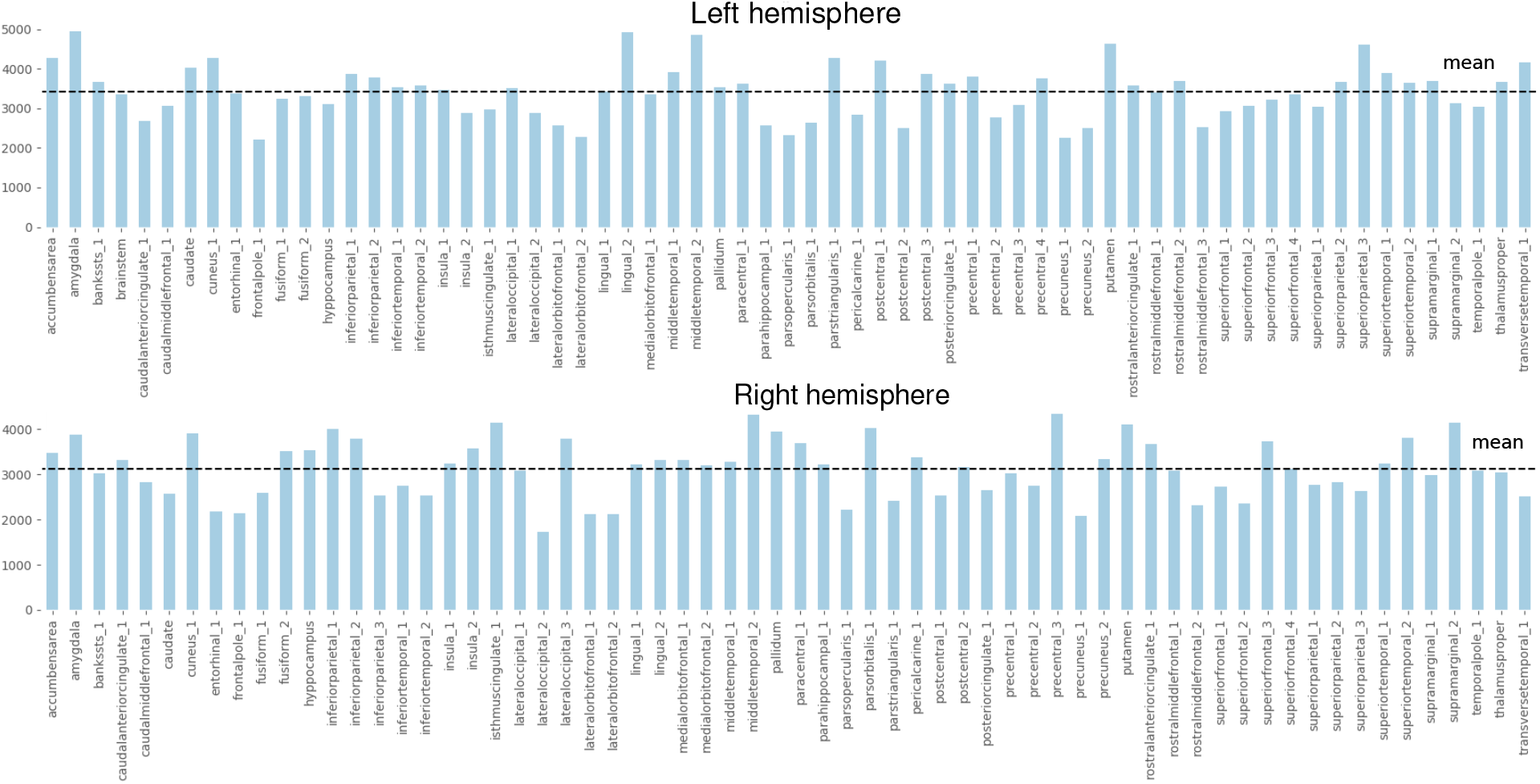
Degree of relevance for ROIs in the **functional** mode of the multimodal 129 × 129 connectome. The horizontal line is the average degree of relevance. ROIs above the mean belong to our a-core.

**Fig. 16.**
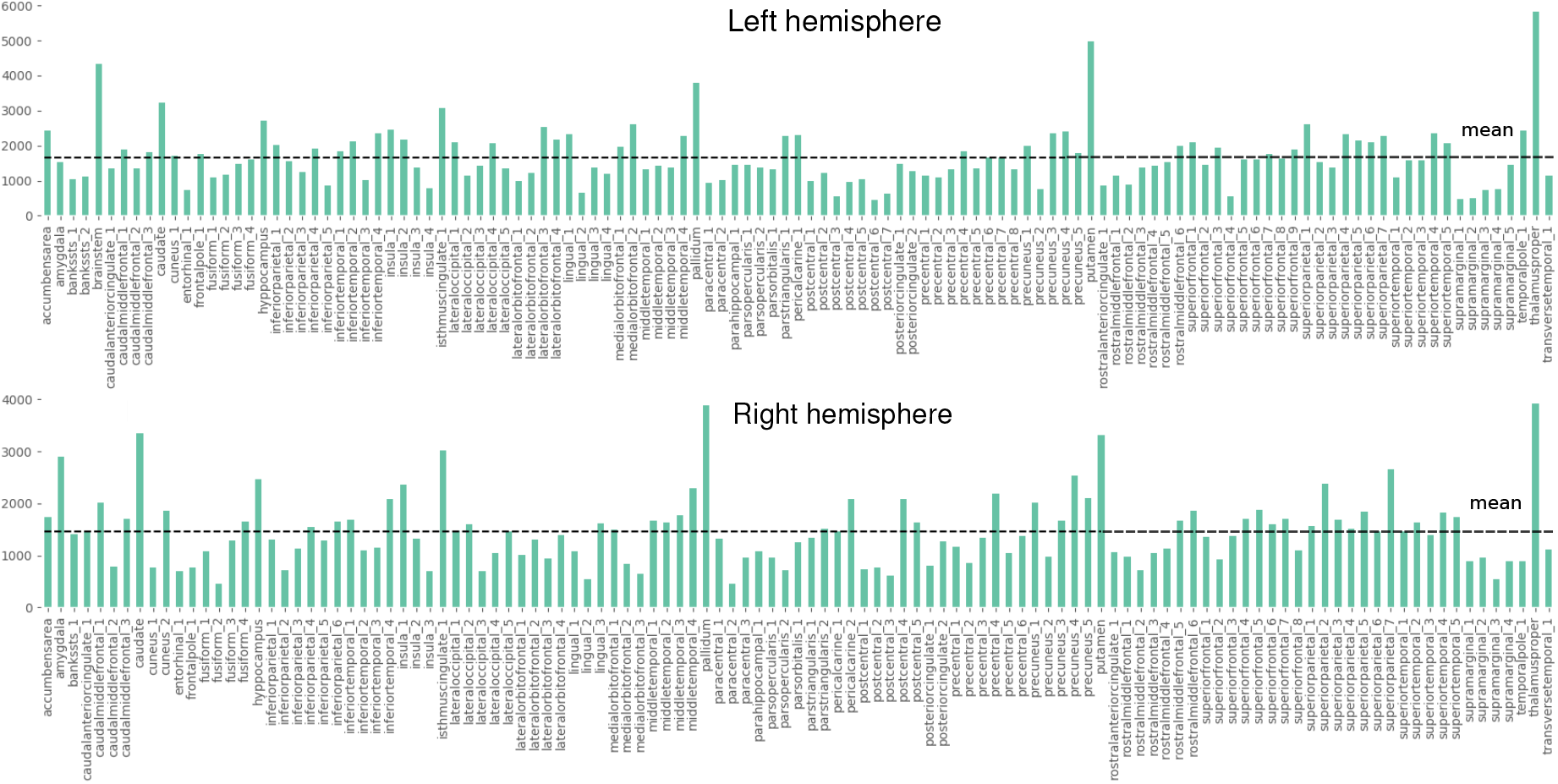
Degree of relevance for ROIs in the **structural** mode of the multimodal 234 × 234 connectome. The horizontal line is the average degree of relevance. ROIs above the mean belong to our a-core.

**Fig. 17.**
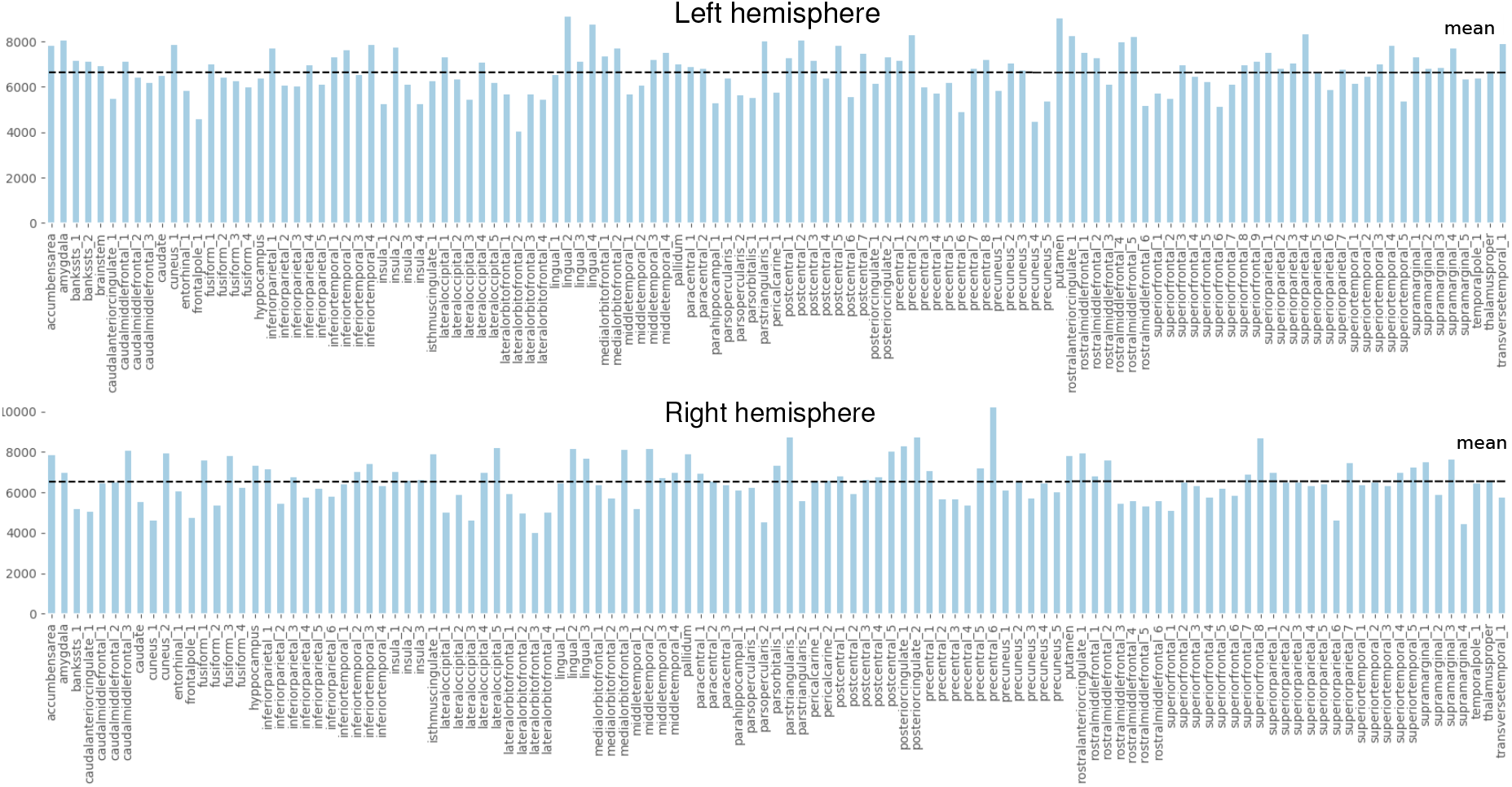
Degree of relevance for ROIs in the **functional** mode of the multimodal 234 × 234 connectome. The horizontal line is the average degree of relevance. ROIs above the mean belong to our a-core.

## Footnotes

1 American Psychiatric Association (2000): Diagnostic and Statistical Manual of Mental Disorders, 4th ed. DSM-IV-TR. American Psychiatric Pub, Arlington, VA22209, USA.

2 https://doi.org/10.5281/zenodo.3758534

3 The source code can be downloaded from https://github.com/leoguti85/BiomarkersSCHZ

